# Alternative designs lead to similar performance when traits and performance vary on different axes

**DOI:** 10.1101/2022.10.05.511045

**Authors:** Kristen M. Nolting, Kent E. Holsinger

## Abstract

Plants differ from one another in size, architecture, water relations, and resource uptake, and these differences often lead to differences in performance. Yet within a community species that differ markedly in these traits often have similar performance. Here we use a simple model to show that when the major axes of trait covariation do not align with the axis of performance variation, large differences among species in structural traits may have similar performance, i.e., ‘alternative designs.’ We further illustrate this phenomenon using trait and performance data from co-occurring *Protea* species in the Cape Floristic Region, South Africa. Long-term coexistence of species within a community requires both similar levels of performance, so that some species are not excluded by competition, and niche differentiation, so that multiple species can coexist. Thus, misalignment between the axis of performance variation and the major axes of trait variation may be common, just as genetic variation may be maintained within a population when the selection gradient does not align with the major axes of the genetic variance-covariance matrix.

## INTRODUCTION

Hutchinson noted in his seminal paper – *“An Homage to Santa Rosalia”* – that there seem to be more species on the planet than niches to hold them (Hutchinson 1959). Although a few authors have argued that trait differences among species do not lead to competitive differences and that species diversity can be largely accounted for by neutral processes (Bell 2000, 2001; Hubbell 2001, 2005; Chave 2004; Alonso *et al*. 2006), most ecologists hold that niche differentiation is required for the maintenance of species diversity within communities (Macarthur & Levins 1967; Chesson 2000; Chase & Leibold 2003; HilleRisLambers *et al*. 2012). Community ecologists often use trait differences among co-occurring species as indices of niche differentiation (Schoener 1968; Brown & Lieberman 1973; Cody 1986; Gillespie 2004; Treurnicht *et al*. 2020), and more recently functional ecologists have identified structural traits, i.e., morphological and allocation-related traits that link an organism’s phenotype with its performance (Violle *et al*. 2007), as important for niche differentiation in diverse community assemblages (Mcgill *et al*. 2006; Cavender-Bares *et al*. 2009; Funk *et al*. 2017).

Global surveys of structural trait variation in land plants reveal large differences among species (Reich *et al*. 1997; Wright *et al*. 2004; Díaz *et al*. 2016), and studies of trait differences among species within diverse clades such as *Helianthus* demonstrate that the magnitude of interspecific trait variation within a clade can rival that seen in global surveys (Mason & Donovan 2015). Even within local plant communities and assemblages traits may differ by orders of magnitude among species (Wright *et al*. 2004; Baraloto *et al*. 2010; Aiello-Lammens *et al*. 2017). The variation within communities may even be greater than expected given regional species pools (Swenson *et al*. 2012). In short, trait variation is abundant at almost any scale, even among individuals within a species growing at a single site (Messier *et al*. 2010; Umaña & Swenson 2019; Nolting *et al*. 2021; Taseski *et al*. 2021). In contrast, substantial performance differences among co-occurring species are less commonly documented (Becker *et al*. 1999; Worthy *et al*. 2020; Nolting *et al*. 2021). Why do we fail to see large differences in performance when we see large differences in structural traits that are related to performance?

Nearly twenty years ago Marks and Lechowicz (2006) hinted at a solution to this paradox. They showed that selection on combinations of traits within a single environment can produce different functional forms (‘alternative designs’) with roughly equivalent performance. Their result suggests that while individual ***trait*** differences among species may reflect differential resource use (and niche partitioning), ***performance*** differences among species within a community are likely to be minimal, a conclusion consistent with coexistence theory (Chesson 2000; Clark *et al*. 2007). Marks and Lechowicz (2006) argue that while trait variation in natural communities is often assumed to reflect species’ optimization to microenvironments, it could be that “…aside from major environmental heterogeneity, alternative designs are a major, if not the main, cause of functional diversity in nature” (Marks & Lechowicz 2006). In spite of their provocative suggestion, few studies have investigated this possibility, and the mechanism that leads to ‘alternative designs’ has not been described.

Here we demonstrate how a particular pattern of trait covariation among ‘alternative designs’ leads to similar performance. First, we use a simple, two trait simulation of performance variation under different scenarios of trait correlations and trait-performance associations, to illustrate that ‘alternative designs’ lead to similar performance only when the major axis of trait variation is not aligned with the axis of performance variation. We then develop a straightforward mathematical model showing that this result holds for an arbitrary number of traits.

We conclude by illustrating the phenomenon in a recently analyzed empirical dataset. Specifically, we show that multivariate combinations of leaf and wood structural traits differ markedly among species but lead to similar physiological performance among co-occurring *Protea s*pecies in the Cape Floristic Region, South Africa (data originally published in Nolting et al. 2021). Taken together, the model and the empirical results suggest the presence of persistent ‘alternative designs’ (i.e., large variation among species in individual structural traits that converge on similar performance, (Marks & Lechowicz 2006; Marks 2007)) within natural communities could be the result of processes that minimize performance differences among species. If ‘alternative designs’ are common, these results have important implications for understanding the ecological and evolutionary mechanisms contributing to persistence of species-diverse assemblages world-wide.

## EXPLORING ALTERNATIVE DESIGNS WITH A SIMPLE MODEL

We begin by developing our intuition with a straightforward model in which individual performance is related to two traits. These traits may be either positively correlated, negatively correlated, or uncorrelated, and the axis along which performance varies may be either aligned or misaligned with the major axis of trait variation.

### Simulating Variance in Performance in a Two-Trait Model

Assume that a single performance trait is linearly predicted by two structural traits:

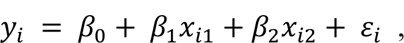

where *y_i_* is the performance of individual *i, x_i1_* and *x_i2_* are the traits of individual *i,* and *ϵ*_i_ is the residual error. *β_1_* and *β_2_* reflect the associations between traits *x_1_* and *x_2_* with *y*, and *β_0_* is the intercept. For simplicity, we will assume that *x_1_* and *x_2_* are centered and scaled to have a mean of zero and standard deviation of 1. Traits *x_1_* and *x_2_* may be correlated, and the correlation coefficient between them is given by *ρ*.

For the positive trait correlation simulations, we assume *ρ* = 0.8 (i.e., a strong positive trait correlation), and we explore two functional axis scenarios: one in which the regression coefficients (*β_1_* and *β_2_*) describing the relationship between our two traits and performance are both positive and of the same magnitude, i.e., the axis of performance variation is aligned with the major axis of trait variation, and a second scenario in which the regression coefficients are of the same magnitude but opposite in sign, i.e., the axis of performance variation is orthogonal to the major axis of trait variation (Figure 1, ‘Positive Trait Correlation’ Box):

**Figure 1.**
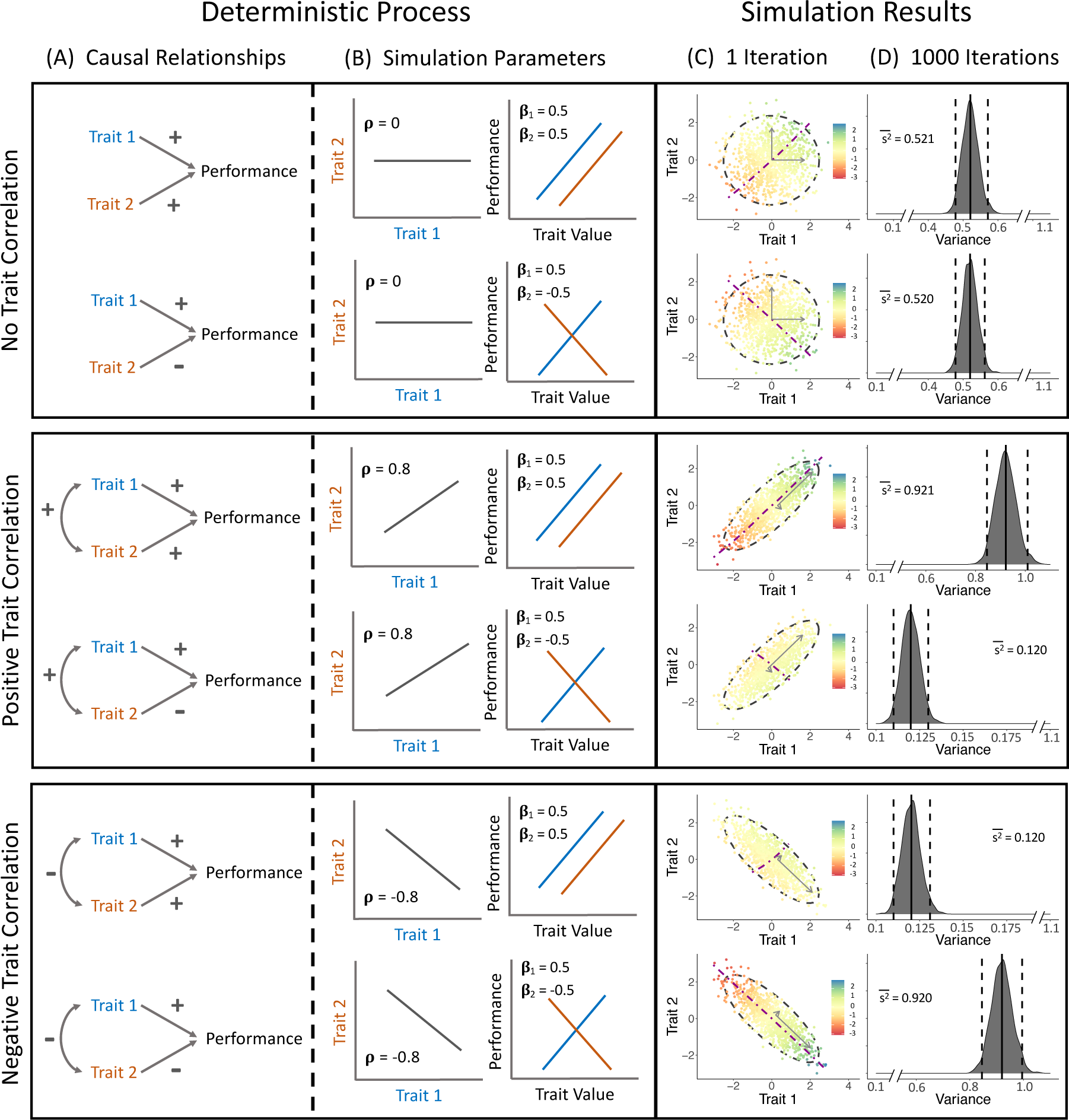
The simple two-trait models and the simulated trait data and distribution of performance variation. Three types of trait correlations are presented: Top Box, no trait correlation between the two traits; Middle Box, a positive trait correlation (*ρ* = 0.8); Bottom Box, a negative trait correlation (*ρ* = −0.8). For each trait correlation, we simulate trait data under two different scenarios: Scenario 1, in which both traits are positively associated with performance (*β*_1_ = 0.5 and *β*_2_ = 0.5), and Scenario 2, in which one trait is positively associated with performance (*β*_1_ = 0.5) and one trait is negatively associated with performance (*β*_2_ = −0.5). For each trait correlation pattern and each scenario, we simulate trait data following the linear models described in the main text. In panel C we present a single realization of the simulation, where the color of each point reflects the predicted performance given the combination of trait values and the linear model defined. The black dashed ellipse around each cloud of points is the 95% confidence interval, the magenta dashed line reflects the functional axis describing the association between the traits and performance, and the gray arrows represent the principal components describing the main axis of trait variation (long arrow) and the minor axis of trait variation (short arrow). We repeated each simulation 1000 times and present the distribution of performance variances under each scenario and trait correlation in panel D. In panel D the dashed vertical black lines reflect the 95% cut-offs for each distribution and the solid black line is the mean variance given 1000 iterations of a simulation.

**Scenario 1:** *β*_1_ = 0.5, *β*_2_ = 0.5.

**Scenario 2:** *β*_1_ = 0.5, *β*_2_ = −0.5.

We also assume *β*_0_ = 0 (i.e., the expected intercept is zero) in our models under each scenario, which is a reasonable approximation if we centered and scaled the response variable as well as the covariates. Finally, we display the outcomes under these two scenarios, by simulating trait data under each scenario for 1000 individuals, sampled from a multivariate normal distribution. The performance value, *y_i_*, for each individual in each scenario was determined from the simulated trait data and the linear model described for each scenario, assuming a residual variance of 0.02. Figure 1c (‘Positive Trait Correlation’ box) reflects the outcome of a single realization of this process, under our two scenarios (scenario 1 above, scenario 2 below). We also report the variance in our performance trait across 1000 realizations for each scenario, and we summarize the distribution of reported variance across realizations in Figure 1d (scenario 1 above, scenario 2 below). For the negative correlation (*ρ* = −0.8) and no correlation (*ρ* = 0) simulations we use the same two functional axis scenarios.

Panel c of Figure 1 shows the distribution of simulated trait data for our three different patterns of trait correlation: no correlation (the top box), positive correlation (the middle box), and negative correlation (the bottom box). Within panel c (Figure 1) for each correlation structure, we show the results from one realization of the trait simulation. The color of each point corresponds to the performance predicted for that individual from the linear regression depending on the scenario chosen, scenario 1 in the top plot and scenario 2 in the bottom plot. The magenta dashed line in each plot represents the regression vector associated with performance in each scenario (what we refer to as the ‘functional axis’ of trait variation), and the gray arrows are the principal components associated with the trait data (with PC1 reflecting the main axis of trait variation and PC2 reflecting the minor axis of trait variation). Panel d of Figure 1 shows the distribution of performance variation from 1000 independent simulations of each scenario and each pattern of trait correlation.

The top box in Figure 1 (‘No Trait Correlation’ box) illustrates that when traits are uncorrelated, the relationship between the major axis of trait variation and the axis of functional variation has no impact on the amount of performance variation observed. In contrast, the middle box (‘Positive Trait Correlation’ box) and bottom box (‘Negative Trait Correlation’ box) both show that when traits are associated with one another the amount of performance variation observed depends on whether the major axis of trait variation is aligned with the axis of functional variation. Specifically, whether traits are positively associated (middle box) or negatively associated (bottom box), there is substantially more performance variation when the functional axis, i.e., the regression vector, is aligned with the major axis of trait variation, PC 1, than when it is aligned with the minor axis of trait variation PC 2. In short, whether we find ‘alternative designs’ depends not on whether traits are negatively correlated – the classic ‘tradeoff’ example often invoked to explain such a phenomenon (Agrawal 2019) – but on ***both*** the pattern of trait correlation ***and*** the pattern of functional variation.

## A GENERAL MODEL FOR ALTERNATIVE DESIGNS

Here we provide a straightforward multivariate extension of the two-trait model we just illustrated:

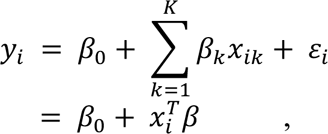

where *y_i_* is the observed physiological performance of individual *i*, *β*_0_ is the intercept, *x_ik_* is the trait value for trait k in individual *i*, *β*_k_ is the regression coefficient for trait k, and *ϵ_i_* is the residual error (with mean 0 and variance σ^2^). *x_i_* is the vector of traits in individual *i* and *β* = (*β*_1,⋯,_*β*_k_), k = 1, . . ., *K* is the regression vector. We assume that traits are drawn independently from a multivariate distribution with mean *μ* and covariance *Σ*_*W*_. We will take expectations with respect to the distribution of *x_ik_*.

The variance of *y* is

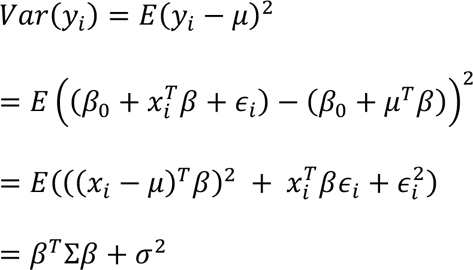

Using the spectral decomposition theorem (Szabo 2015) we can rewrite Σ as

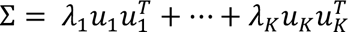

where the eigenvalues *λ_i_* are numbered from the largest to the smallest, and the corresponding eigenvalues have unit length. This allows us to rewrite *Var*(*y_i_*) as

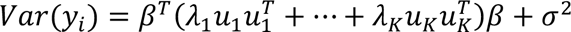

Suppose that the vector of regression coefficients is parallel with the *i*th principal component. It is, by definition, orthogonal to all of the remaining components, and *Var*(*y_i_*) reduces to

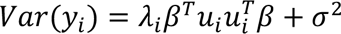

If the *i*th principal component is one of the minor axes of trait variation, the variation in performance will be much smaller than if it is parallel to one of the major axes of trait variation. More generally, if the regression vector, i.e., the functional axis along which individuals differ in performance, is mostly orthogonal to the major axes of trait variation, there will be little variation in performance.

## EXAMPLE: PROTEAS WITH DIFFERENT TRAITS HAVE SIMILAR PERFORMANCE

### Alignment of the Functional Axis with Axes of Trait Variation

The models above show that ‘alternative designs’ may arise when the major axes of trait variation do not align with the axis of functional variation. To determine whether we can detect this phenomenon in empirical data, we re-examine a subset of a previously published dataset that measured many structural traits (e.g., leaf area, wood density, stomatal size) and several aspects of physiological performance (e.g., photosynthetic rate, stem hydraulic conductance) on naturally co-occurring *Protea* species in the Cape Floristic Region in southwestern South Africa (Nolting et al. 2021; see supplement for additional information about the species included and trait sampling). Nolting et al. (2021) reported that (a) combinations of structural trait values explain much of the variance in physiological performance traits (here our full model R^2^ values range from 0.26 to 0.44, Table 1; Figure S1), (b) species differ markedly in structural traits (with estimated among-species variation in structural traits ranging from 54.9 to 95.1%; Figure S2a), but (c) species largely overlap with respect to physiological performance traits (estimated among-species variation in performance traits ranging from 3.2 to 31.5%; Table 1; Figure S2b). See supplemental materials for more information about model fits and variance partitioning.

**Table 1.** Model fits and estimated among-species variance for each physiological performance response trait explored in the *Protea* dataset. The mean estimated Bayesian R^2^ for the full model and for the fixed effects only (the marginal R^2^) are presented for each model, as well as the mean estimated among species variance for each physiological performance trait, relative to the residual variance (i.e., within species variance and residual error).

| Physiological<br>Performance Measure | Full Model $R^2$<br>(Marginal $R^2$ in parentheses) | Estimated Among Species<br>Variance (%) |
| --- | --- | --- |
| Sapwood-Specific Hydraulic<br>Conductivity | 0.36 (0.20) | 27.2 |
| Leaf-Specific Conductivity | 0.30 (0.22) | 26.6 |
| Photosynthetic Rate | 0.44 (0.27) | 31.5 |
| Leaf-Specific Photosynthetic<br>Rate | 0.33 (0.36) | 18.0 |
| Stomatal Conductance | 0.36 (0.18) | 7.3 |
| Instantaneous Water-Use<br>Efficiency | 0.26 (0.20) | 3.2 |

Using the princomp function in R (R Core Team 2022; version 4.2.1) we performed a principal components analysis on the centered and scaled structural trait data in our *Protea* dataset, yielding eight principal components. Then for each of the six physiological performance measures we evaluated the alignment of the structural trait PC axes with the vector of regression coefficients describing the association between traits and physiological performance, i.e., the functional axis. Nolting et al. (2021) included site and species random effects in their analyses of trait-performance relationships, but we exclude them here to facilitate comparison of the empirical results with the model expectations developed above.

We quantified the degree of alignment for each performance trait model by calculating the angle between the vector of regression coefficients for each response and each of the eight principal components (Figure 2). An angle close to 90 degrees indicates that the functional axis is almost completely misaligned with that principal component axis. An angle close to zero degrees indicates that the functional axis is almost completely aligned with that principal component (Figure 2). While the degree to which principal component axes align with the functional axis differs among physiological performance traits, for every performance trait the major axes of trait variation (i.e., PC axes 1 and 2) are generally the axes most misaligned with the functional axis. Instead, the minor axes of trait variation (often the axes excluded from analyses that use dimension reduction to estimate functional associations from trait matrices and performance) are the axes most parallel with the regression vector reflecting the functional axis.

**Figure 2:**
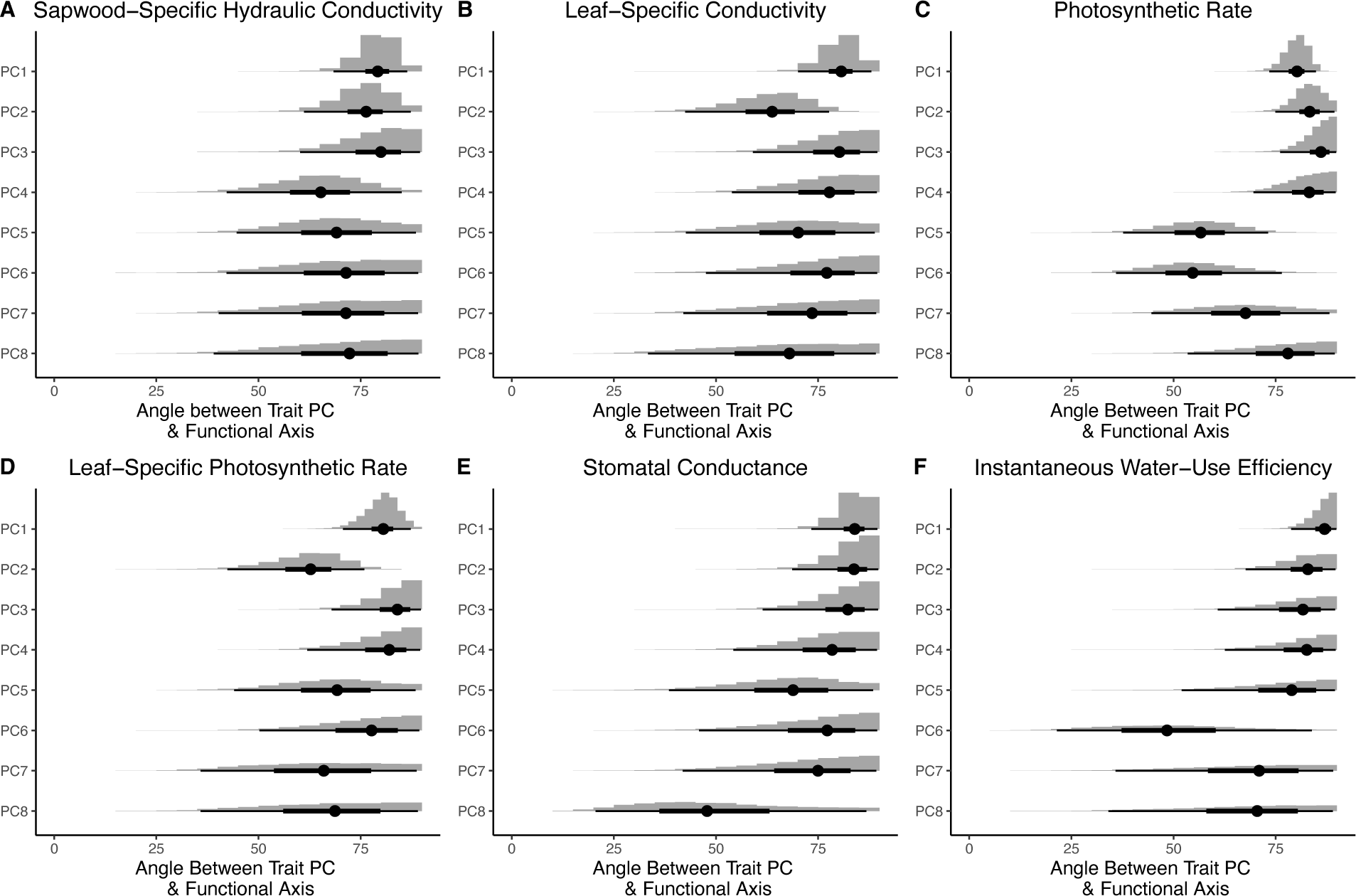
Alignment of each principal component with the regression vector describing the functional relationship between structural traits and each of six performance traits in the *Protea* dataset. (A) Sapwood-specific conductivity, (B) Leaf-specific conductivity, (C) Photosynthetic rate, (D) Leaf-specific photosynthetic rate, (E) Stomatal conductance, and (F) Instantaneous water-use efficiency. The horizontal axis is the angle (0-90°) between each of the eight principal components for the structural trait data measured on 132 *Protea* individuals and the regression vector associated with each physiological performance trait. Angles closer to 90° indicate that the functional axis (i.e., the vector of regression coefficients) and the PC axis are largely orthogonal and angles closer to 0° indicate they are more aligned. The gray histograms reflect the posterior distribution of angles, the black circles represent the mean of the posterior, the thick black lines reflect the 50 percent credible intervals, and the thin black line reflects the 95 percent credible intervals.

We see little differentiation in physiological performance among *Protea* species despite considerable trait differences because the axis along which trait differences are associated with differences in physiological performance is largely orthogonal to the axes that account for most structural trait variation. An important implication of this result is that reducing trait variation to the first few PCs and attempting to associate those with functional differences may fail to find real associations of traits with functional differences.

### Different Structural Trait Combinations Lead to Similar Performance within and across Species

Our results show that when the axis of performance variation does not align with the major axes of trait variation, different trait combinations may lead to similar measures of performance. Moreover, these ‘alternative designs’ (Marks & Lechowicz 2006; Marks 2007) might exist not only across taxa but also *within* species that demonstrate substantial trait variation.

To illustrate how combinations of structural traits can lead to similar performance both within and across species in our dataset, we plot our sampled *Protea* individuals in multivariate structural trait space, but color the individuals based on their relative performance values. Each point in each panel in the top row of Figure 3 (Figure 3a,b,c: using the first two PC axes) reflect the position of individuals in the first two dimensions of structural trait space, i.e., the two most important axes of structural trait variation. In each panel the points are in the same position, but their coloring reflects the relative performance of each individual for one of three physiological performance traits. For example, in Figure 3a the ***position*** of each sampled individual is determined by its position in the structural trait PCA, but the ***coloring*** of each individual is based on the quintile of K_s_ (sapwood-specific hydraulic conductivity) into which it falls. Confidence ellipses in this top panel of Figure 3 (Figure 3a,b,c) delimit the traits within a 95 percent confidence interval of each species mean. Notice that in the part of trait space accounting for most variation, species are well differentiated from one another. In the bottom panel of Figure 3 (Figure 3d,e,f), we color the points in the same way but plot individuals in structural trait space using the PCs that are most aligned with performance (i.e., the two PCs that were closest to parallel with the regression vector, from Figure 2). Notice that in the part of trait space most closely aligned with performance, species boundaries overlap broadly.

**Figure 3:**
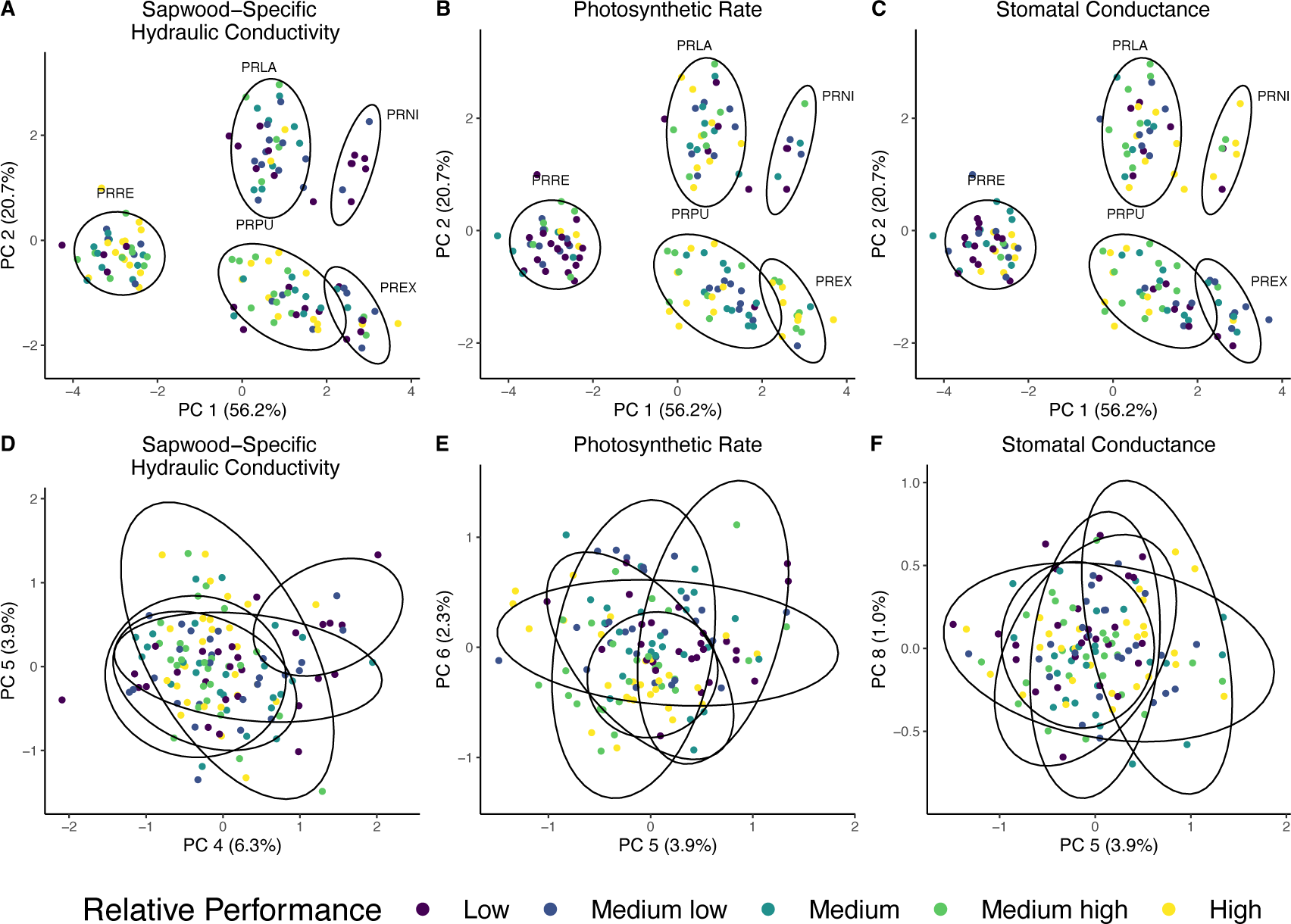
Visualizing variation in relative performance arising from different combinations of structural traits, for a subset of performance traits (Sapwood-specific hydraulic conductivity: A and D; Photosynthetic rate: B and E; Stomatal conductance: C and F). For each panel, individuals from our empirical dataset are plotted with respect to their position in multivariate structural trait space. For A, B, and C individuals are plotted using the first two principal components (PC1 and PC2). For D, E, and F individuals are plotted in the structural trait space that is most aligned with performance (i.e., the PC axes that are most parallel with the regression vector for each performance trait, from Figure 2). The color of the points reflects the relative performance of each individual, corresponding to one of five ordered categories (Low, Medium Low, Medium, Medium High, and High). The confidence ellipses correspond to species groupings. PRLA: *P. laurifolia*, PRNI: *P. nitida*, PREX: *P. eximia*, PRPU: *P. punctata*, and PRRE: *P. repens*.

Closer examination of Figure 3 also shows both that individuals with different combinations of structural traits (i.e., different positions in PCA space) can have similar performance (i.e., fall into the same performance quintile - the same color) and that many combinations of traits lead to performance in the highest (or lowest) quintile. In addition, not only may individuals within species fall into different performance quintiles, but individuals belonging to different species may fall into the same performance quintile. In fact, all species (other than *P*. *nitida*) have some individuals with sapwood-specific hydraulic conductivity, K_s_, in the highest performance quintile and others in the lowest performance quintile. The same pattern is repeated across all six physiological performance traits measured (Figure S3).

## DISCUSSION

Given the well-established relationships between plant traits and plant performance, the magnitude of species differences in performance is often much less than might be expected. We demonstrate that this outcome is expected when the axis of functional variation does not align with the major axes of trait variation. Using a recently published dataset of plant structural traits and physiological performance traits measured in several natural populations of *Protea* from the Cape Floristic Region, South Africa (Nolting et al. 2021), we show that large among-species differences in structural traits related to plant performance do not lead to noticeable differences in physiological performance. Consistent with our mathematical model, we show that the axis of observed functional variation is not aligned with the major axes of observed trait variation. These results illustrate that a full understanding of multivariate trait associations requires not only that we evaluate axes of trait variation, but also that we determine whether major axes of trait variation align with axes of performance variation.

### Implications for Trait-Mediated Species Coexistence and Community Ecology

Plants or animals that live in the same place typically experience roughly the same macroenvironment. As a result, we might expect them to have similar traits (Keddy 1992), but basic principles of community ecology suggest that species can coexist long term only if they differ along at least one niche axis (Chesson 2000; HilleRisLambers *et al*. 2012). To put it another way, environmental filtering may explain an association between community-weighted mean trait values and environments (Ackerly & Cornwell 2007) and phylogenetic underdispersion in local community assemblages (Webb 2000; Webb *et al*. 2002; Mayfield & Levine 2010), but within-community trait differences may promote coexistence (Cavender-Bares *et al*. 2004a, b; Kraft *et al*. 2008; Swenson & Enquist 2009).

Marks and Lechowicz (2006) illustrated that ‘alternative designs’ may lead to roughly equivalent functional performance and may resolve this apparent contradiction. As we show in greater detail, many different trait combinations can produce similar performance. This suggests trait differences promote coexistence through niche differentiation, but if the axis of functional diversity is largely orthogonal to the axis of trait differentiation then performance differences are minimized, an observation that has been observed in previous empirical systems (Adler *et al*. 2007; Umaña *et al*. 2016). Given that many structural traits not only influence differences in any one performance trait, but that any one structural trait influences differences in many performance traits (Worthy *et al*. 2020; Funk *et al*. 2021), the trait space where performance is roughly equivalent may be extremely large (Marks & Lechowicz 2006; Laughlin 2014; Laughlin & Messier 2015).

### Should ‘Alternative Designs’ be Common?

Although we provide only one empirical example, we suspect that ‘alternative designs’ are common in natural systems. Here we focused on structural trait and physiological performance trait variation at the level of individual plants and plant communities, but similar phenomena arise in quantitative genetics. It has long been known that in spite of strong stabilizing selection, individual traits under selection can retain substantial genetic variation (Arnold 1981; Falconer 1981; Mousseau & Roff 1987). This putative inconsistency can be understood by considering the complex interaction of selection operating on many covarying traits (Walsh & Blows 2009). Lande and Arnold (1983) showed how to use the multivariate breeder’s equation, Δ**z** = **G*β***, to identify the selection gradient on traits in a population. The selection gradient, ***β***, is the set of regression coefficients relating trait values to fitness – the direct analog of the functional axis of trait variation we have been discussing. Just as we demonstrated above, Walsh and Blows (2009) show that if the selection gradient is not aligned with major axes of the G matrix (**G**, the genetic variance-covariance matrix), then there may be substantial genetic variation for traits that are under stabilizing selection.

Far from being an exception, misalignment between multivariate axes of genetic variation and the selection gradient may actually be expected. It is often thought that the direction of population divergence will correspond with ‘genetic lines of least resistance’, i.e., divergence along the major axes of genetic variation (Schluter 1996), and there have been many examples of this phenomenon presented (Mitchell-Olds 1996; Chenoweth *et al*. 2010). Recent work in *Anolis* lizards suggests that the alignment between major axes of genetic variation and axes of divergence can be maintained over millions of years, and despite evolution of the G matrix itself (McGlothlin *et al*. 2018). Nonetheless, there are many other examples in which divergence seems to be constrained due to misalignment between the major axes of genetic variation and the selection gradient either within or among populations (Etterson & Shaw 2001; Henry & Stinchcombe 2023). Taken together these results indicate that an evolutionary response might be limited not only if a population lacks genetic variation, but also if the selection gradient is not aligned with major axes of the G matrix (Donovan *et al*. 2011; Agrawal 2019). Interestingly, Engen and Saether (2021) suggest that the evolutionary process itself may lead to misalignment of the selection gradient and major axes of the G matrix. Specifically, they show that at an evolutionary equilibrium the G matrix itself evolves so that the selection gradient is parallel to the eigenvector associated with the smallest eigenvalue, i.e., the axis of the G matrix associated with the least variation. We conjecture that the process of interspecific competition will lead to a similar misalignment between the axis of functional variation and the major axes of structural trait variation in natural communities.

## CONCLUSION

Functional ecologists have often focused on broad-scale geographic patterns in which trait values are seen as characteristic of, and perhaps even diagnostic of, particular environments (Diaz *et al*. 1998; Cavender-Bares *et al*. 2004b; Cornwell & Ackerly 2009; Anderegg *et al*. 2021). Community ecologists, on the other hand, have typically focused on trait differences as key features that allow long-term coexistence (Lack 1947; Cody 1986; Silvertown 2004). When these perspectives are combined, we find that there are often large differences between species in structural traits but that performance differences among co-occurring species are minimal, even when those structural traits are related to performance. Marks and Lechowicz (2006) demonstrated that ‘alternative designs’ can evolve within a homogenous environment, and they suggested that ‘alternative designs’ may be a major cause of among-species trait variation observed in plant communities (Marks and Lechowicz, 2006). Here we show how this pattern can emerge. When the major axes of trait variation are largely orthogonal to the functional axis describing the association between traits and performance, there will be little variation in functional performance. Because long-term co-existence among species requires similar levels of performance within a given habitat, we expect the axis of functional performance will often be orthogonal to major axes of trait variation.

The pattern we describe is directly analogous to phenomena that are well described in quantitative genetics. Just as the response to selection is largely the outcome of the alignment between major axes of the G matrix and the selection gradient, so the extent of performance variation across species depends on whether the major axes of trait variation are aligned with the axis of functional variation. While it is tempting to describe ‘alternative designs’ as an example of ‘trade-offs’, i.e., negative covariance between two traits important for performance, we show that ‘alternative designs’ can arise even when traits positively covary. What matters is not whether traits covary positively or negatively, but whether the axis of trait variation is aligned with the axis of functional variation.

## STATEMENT OF AUTHORSHIP

K.M.N. and K.E.H. conceived the study. K.E.H. developed the generalized model with input from K.M.N. K.M.N. performed all analyses, simulations, and generated all figures with input from K.E.H. K.M.N. and K.E.H. developed the biological interpretation of the simulated, mathematical, and empirical results. K.M.N. wrote the first manuscript draft with input from K.E.H.

## DATA AND CODE AVAILABILITY

The data, R scripts, and model fit files underlying all analyses, figures, and results have been deposited in a ZENODO repository (10.5281/zenodo.8083856).

## Supporting information

Supplemental Methods and Results

## ACKNOWLEDGEMENTS

We are very grateful to R. Bagchi who initially gave us the idea to explore this question with respect to trait variances and covariances. We also thank C. Jones, R. Bagchi, and members of the Donovan and Burke Labs for very helpful feedback on earlier versions of this manuscript. This work was supported in part by the National Science Foundation (DEB-1046328).

