## Supplemental Methods and Results for "Alternative designs lead to similar performance when traits and performance vary on different axes"

**This file contains:**

Section 1: additional detail about trait sampling from the empirical *Protea* dataset

Section 2: additional detail about data analysis for the *Protea* dataset

Section 3: additional figures

#### I. EMPIRICAL DATA COLLECTION

We use data from (Nolting *et al.* 2021) to explore the relationship between structural trait variation and physiological performance. Specifically, we demonstrate that (1) traits predict physiological performance relatively well, and (2) species differ markedly in their traits, but do not substantially differ in their physiological performance.

*Study Site and Species*

Nolting et al. (2021) investigated multivariate trait-performance relationships for five *Protea* species sampled across three sites in the Cape Floristic Region, South Africa. The Cape Floristic Region (CFR) is the smallest of the world's six floral kingdoms (area ~90,000 km$^2$), but it hosts over 9,000 plant species (Linder 2003; Peter Linder 2005) making it the most diverse of the five Mediterranean regions (Cowling *et al.* 1996). Like other Mediterranean biomes, the CFR is characterized by warm, dry summers and cool, wet winters (Cowling *et al.* 1996), but there is also a west-east seasonality gradient with winter rainfall in the west and relatively constant, low rainfall in the east. *Protea* species primarily occur in fynbos habitat, which is characterized as a shrubland with nutrient-poor soils (Allsopp, N. Colville, J.F. Verboom, G.A. 2014) and natural, recurring fires that sweep through sites on average every 10-12 years (Van Wilgen *et al.* 2010).

The genus *Protea* L. (Proteaceae) represents a morphologically diverse (~112 species in Africa, with ~60% endemic to the Cape Floristic Region; (Schnitzler *et al.* 2011)), iconic, and locally abundant plant genus that typifies many fynbos communities. *Protea* exhibit a diversity of form and function with respect to growth form (e.g., low growing, prostrate shrubs to upright shrubby trees), response to fire (e.g., reseeding species that die with fire but release all of their seeds during the fire event from persistent infructescences versus resprouting species that regrow after fire in addition to releasing seeds), and vegetative traits associated with carbon and water use strategies (e.g., leaf size and shape variation). Previous studies on *Protea* have demonstrated that variation in structural traits (e.g., leaf mass per area and wood density) and physiological traits (e.g., photosynthetic rates and stem hydraulic conductivity) are associated with fitness differences (Carlson *et al.* 2011; Prunier *et al.* 2012; Nolting *et al.* 2021). Further, variation in these traits is associated with variation in the abiotic environment, such as precipitation and temperature gradients across the CFR (Carlson *et al.* 2011; Mitchell *et al.* 2015).

*Trait Collection*

The samples included in Nolting et al. (2021) occur at three sites and include 11-20 individuals of each species per site (a total of 9 populations). Due to complexity in trait distributions, we excluded one population (*Protea repens* at Cederberg) in our reanalysis here in order to simplify the presentation of the empirical data with respect to the mathematical model presented. This leaves us a total of 8 populations and 132 sampled individuals. On each individual plant sampled, Nolting et al. (2021) recorded measures of plant size (e.g., height and canopy area) and reproductive effort measured as the total number of inflorescences and infructescences retained on a plant. The *Protea* species included in the study either regenerate from seed or resprout after fire, meaning that adults (a) are all roughly the same age within a stand as they either recruit as seedlings together post-fire or regrow from persistent stems, and (b) all retain their infructescences until these ‘seedheads’ split and release seeds during a fire. These characteristics make it possible to record reproductive effort for individuals (e.g., inflorescences plus infructescences) which has been shown to covary with seed production (Carlson & Holsinger 2013), representing a proxy for fitness (Nolting *et al.* 2021).

On each individual, Nolting et al. (2021) also measured area-based, light-saturated photosynthetic rates ($A_{area}$), stomatal conductance ($g_s$), and instantaneous water-use efficiency ($WUE_{Instan}$) in the field. They collected branches to measure sapwood-specific hydraulic conductivity ($K_s$) following Espino and Schenk (2011), and quantified the Huber value of each stem (the sapwood cross-sectional area of a stem relative to the total one-sided leaf area per area, (Mencuccini & Bonosi 2001)) on which hydraulic measures were recorded, to calculate leaf-specific conductivity (LSC, sapwood-specific conductivity divided by total leaf area of the stem, Tyree and Zimmerman 2002) and leaf-specific photosynthetic rates (LSP, light-saturated photosynthetic rate per area divided by Huber value). From the stems and leaves on which physiological traits were measured, Nolting et al. (2021) measured a suite of structural traits (e.g., traits related to organ morphology and allocation): leaf area (LA, $cm^2$), leaf mass per area (LMA, g $cm^{-2}$), lamina density (LD, g $cm^{-3}$), leaf length-width ratio (LWR, unitless), stomatal pore length (SL, mm), stomatal pore density (SD, $mm^{-2}$), bark thickness (BT, mm), and wood density (WD, g $cm^{-3}$). These structural and physiological performance traits were chosen based on the large literature in plant functional ecology suggesting that they are key indices of carbon uptake and allocation and water-use strategies (see Nolting et al. 2021 for further discussion of functional significance of each trait, and of trait complexes). Further, Nolting et al. (2021) demonstrate that many of these structural and physiological performance traits are directly and/or indirectly associated with individual plant size and reproductive effort.

#### II. PROTEA DATA ANALYSIS

Nolting et al. (2021) showed that the measures of physiological performance we investigate below are associated with plant size and reproductive effort, which we take to be reasonable proxies for fitness. Thus, we focus on investigating the relationship between eight individual structural traits and six measures of physiological performance.

*Structural Traits Predict Physiological Performance*

We evaluated the relationship between structural traits and each of our physiological performance traits using Bayesian linear mixed effects models (one model for each of the six physiological performance traits) using the package 'brms' (Bürkner 2017, 2018, 2021) (version 2.18.0) in the R programing environment (R Core Team 2022; version 4.2.1). All performance and structural traits were standardized to a mean of zero and a unit standard deviation prior to analysis. Sample site, species identity, and the interaction between site and species were included as random effects. We used independent N(0,1) priors on the intercept and regression coefficients, and half Cauchy (0,5) priors on the variance parameters. We used four chains with a warm-up of 2000 iterations and a sampling of 2000 iterations for a total of 8000 samples from the posterior with delta = 0.999. All parameters had Rhat < 1.01, and no divergent transitions were reported. In Figure S1 we plot the predicted performance values versus the observed values, and report the mean Bayesian $R^2$ estimates (Gelman et al. 2017). Consistent with Nolting et al. (2021), we demonstrate that combinations of structural traits predict performance reasonably well (i.e., full model $R^2$ values range from 0.26 to 0.44; marginal $R^2$ values – fixed effects only – ranging from 0.18 to 0.36).

*Species Differ in Structural Traits but Not in Physiological Performance Traits*

We evaluated the among species variation (relative to residual variation) in each of our structural and performance traits using a simple regression model with species as a random intercept. We fit these models in 'brms' as above, using the same priors, model specification, and sampling. As Nolting et al. (2021) discuss in more detail, the proportion of variation among species in structural traits (Figure S2a) is much greater than that in physiological performance traits (Figure S2b). As individuals sampled were of similar size within each population, size-mediated trait differences are unlikely to contribute largely to the observed differences. In short, there are substantial differences among species in structural traits that are associated with performance, but there are only minimal differences among species in physiological performance traits.

### III. ADDITIONAL FIGURES

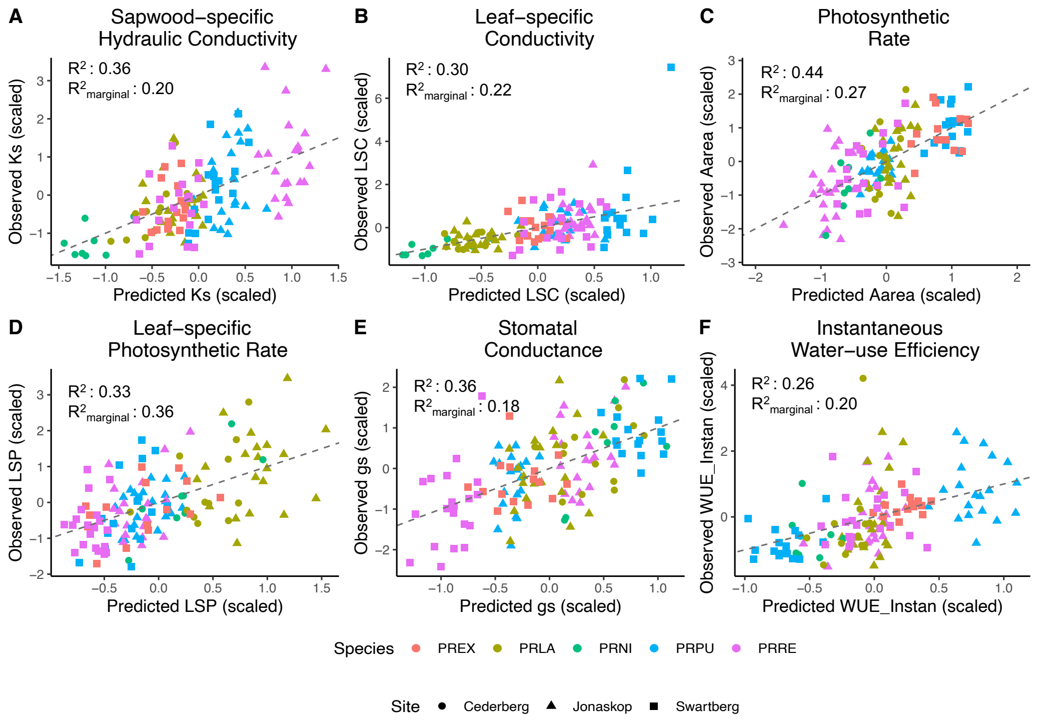


**Figure S1:** For each multiple regression modeling the relationship between the eight structural traits and each of the six performance traits, we plot the predicted performance values versus observed (with traits centered with a mean of zero and scaled to unit variance). (A) Ks: Stem-specific hydraulic conductivity, (B) LSC: Leaf-specific conductivity, (C) $A_{area}$: Photosynthetic rate, (D) LSP: Leaf-specific photosynthetic rate, (E) $g_s$: Stomatal conductance, and (F) WUE: Instantaneous water-use efficiency. The dotted gray line reflects the 1:1 line. Colors represent different species and shapes represent different sites. We report the mean Bayesian $R^2$ value for the full model and for the fixed effects only (i.e., marginal $R^2$), for each model.

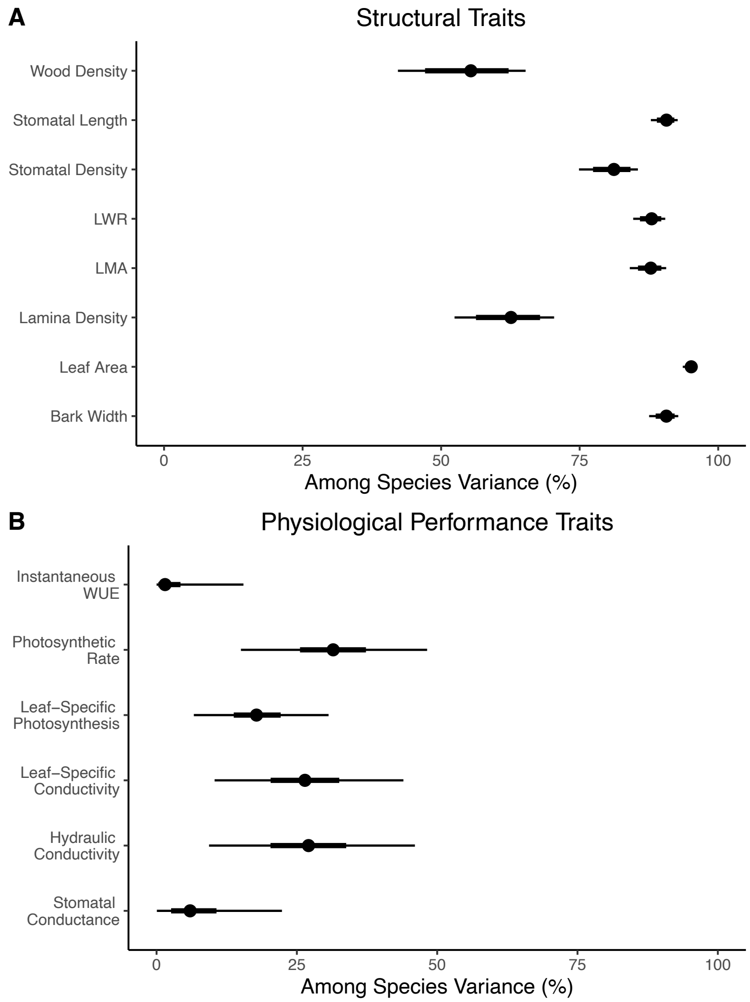


**Figure S2:** Among species variance partitioning for the eight structural traits (A) and the six performance traits (B). The gray histograms reflect the full posterior distribution, the black circles represent the mean of the posterior, the thick black lines reflect the 50 percent credible intervals, and the thin black line reflects the 95 percent credible intervals. Adapted from Nolting et al. 2021, with permission from *Annals of Botany* pending.

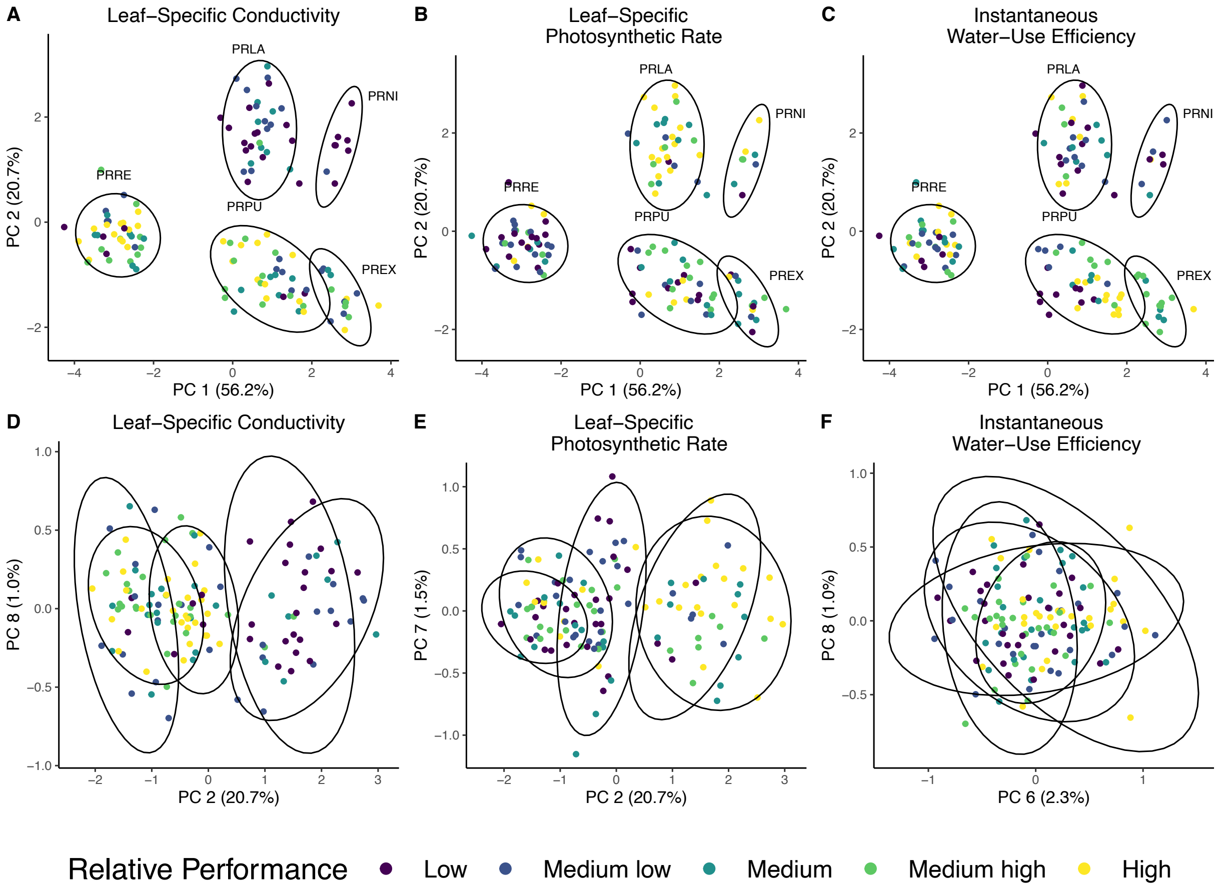


**Figure S3:** Visualizing variation in relative performance arising from different combinations of structural traits, for a subset of performance traits (Leaf-specific conductivity: A and D; Leaf-specific photosynthetic rate: B and E; Instantaneous water-use efficiency: C and F). For each panel, individuals from our empirical dataset are plotted with respect to their position in multivariate structural trait space. For A, B, and C individuals are plotted using the first two principal components (PC1 and PC2). For D, E, and F individuals are plotted in the structural trait space that is most aligned with performance (i.e., the PC axes that are most parallel with the regression vector for each performance trait). The color of the points reflects the relative performance of each individual, corresponding to one of five ordered categories (Low, Medium Low, Medium, Medium High, and High). The confidence ellipses correspond to species groupings. PRLA: *P. laurifolia*, PRNI: *P. nitida*, PREX: *P. eximia*, PRPU: *P. punctata*, and PRRE: *P. repens*.
